# An Integrative Taste Receptor Links pH and Amino Acids to Sugar Sensing in *Bemisia tabaci*

**DOI:** 10.64898/2026.05.28.728435

**Authors:** Ofer Aidlin Harari, Esther Yakir, Victor Santos, Dor Wintraube, Ella Tadmor, Ksenia Juravel, Karmit Levy, Hodaya Glik, Jonathan D. Bohbot, Shai Morin, Osnat Malka

**Affiliations:** Department of Entomology, Institute of Environmental Sciences, The Robert H. Smith Faculty of Agriculture, Food and Environment, The Hebrew University of Jerusalem, Rehovot, Israel

## Abstract

Phloem-feeding insects execute complex behavioral decisions to secure essential nutrients from a diet characterized by nitrogen scarcity and severe osmotic pressure due to high sucrose concentrations. We investigated the sensory mechanisms underlying these decisions in the phloem-feeding whitefly *Bemisia tabaci*. We demonstrate that the sweet taste receptor BtabGR1, expressed in mouthpart and gut tissues, integrates three environmental chemical cues: sucrose concentration, the presence of the essential amino acid arginine, and pH values. Arginine is a pH-dependent positive modulator of sucrose sensing, increasing receptor responses nearly fourfold under apoplast-like conditions and more than doubling the receptor responses in the gut luminal environment. Insects show a strong feeding preference for arginine-containing diets in dual-choice bioassays, with markedly higher intake when arginine is present. RNAi-mediated silencing of *BtabGR1* disrupt intake regulation, leading to increased honeydew excretion. These findings suggest a putative link between arginine and the BtabGR1 receptor in regulating both feeding-site evaluation and diet ingestion. Furthermore, the ability to integrate three distinct environmental cues makes BtabGR1 one of the most complex interdependent sensory systems described for a single insect chemoreceptor.

## Introduction

The phloem vascular tissue is the main pathway for long-distance transport of assimilates from source to sink organs in plants ^1^. As a result, the chemical composition of phloem-sap is unique compared to its surrounding tissues. The dominant components of phloem-sap are sugars, primarily photosynthetically derived sucrose, which is loaded into the phloem^2^ at concentrations ranging from 251 to 1384 mM^3–6^. Amino acids are the second most abundant components, with total concentrations ranging from 40 to 500 mM. The composition of phloem amino acids varies among plant species. However, the non-essential amino acids, glutamate, glutamine, aspartate, asparagine and serine, typically dominate, whereas essential amino acids such as arginine, histidine, methionine and tryptophan occur at lower levels^3,4,7^.

Amino acids are synthesized in mesophyll cells or transported from roots via the xylem^7^, and like sucrose, are subsequently loaded into the phloem for translocation from source (leaf) to sink tissues^8,9^. Accordingly, whereas individual amino acid concentrations in the leaf apoplast (extracellular space) range from undetectable amounts to 0.5 mM (1.7-2.4 mM total), their median concentrations in the phloem are 61-141-fold higher, depending on amino acid identity and plant species^10–12^ (Table S1). Another key distinction between phloem-sap and the surrounding apoplastic fluid is pH. Phloem-sap, like the cytoplasm, is slightly alkaline, with an average pH of ∼8 reported across studies^13–16^. In contrast, the leaf apoplast is generally acidic, with pH values ranging from 4.7 to 6.5 (average ∼5.3), depending on plant species, light conditions, and stress factors^17–22^.

Despite its relatively nutrient-rich and toxin-poor composition compared to other plant tissues, the phloem-sap is exploited as a primary food source only by certain insect lineages within the order Hemiptera^23^. This feeding strategy occurs in some planthoppers (Auchenorrhyncha), some leafhoppers (Clypeorrhyncha), and most sternorrhynchan species, including whiteflies, aphids, mealybugs, and some psyllids^23–25^. Adopting a phloem-feeding lifestyle requires adaptations to several inherent challenges. The sieve-tubes are deeply embedded within plant tissues, making them difficult to locate and access. In addition, phloem-sap presents a highly imbalanced nutrient profile: it is extremely rich in sugar (primarily sucrose), imposing strong osmotic pressure in the gut, while essential amino acids are comparatively scarce. Overall nitrogen levels and the ratio of free amino-acids to sucrose are low relative to other plant tissues, such as young leaves and reproductive organs^23,26–30^.

To overcome the challenge of sieve-tubes access, phloem-feeding insects have evolved specialized needle-like mouthparts or stylets, that penetrate the plant cuticle, epidermis, and mesophyll to reach the phloem sieve-elements. Aphids are thought to follow a preprogrammed “rejection–acceptance” behavioral strategy, in which stylets advance through the apoplast while making periodic intracellular punctures, sampling along the way until suitable sucrose concentrations and pH are detected^31,32^. In a previous study^28^, we identified a molecular mechanism in the whitefly *Bemisia tabaci* that enables tracking of increasing sucrose concentrations from the leaf surface to the sieve-tube via a highly specific and sensitive sucrose receptor, BtabGR1^28^.

To cope with the scarcity of essential amino acids, sternorrhynchan insects such as *B. tabaci* harbor bacterial symbionts within specialized cells called bacteriocytes. These symbionts can synthesize the full complement of essential and non-essential amino acids^27^. Nevertheless, the insects must acquire sufficient nitrogen from their relatively nitrogen-poor diet. To do so, phloem feeders exhibit compensatory feeding, ingesting large volumes of sap to meet their nitrogen requirements^33–35^. The strategy comes at a cost: the high sugar load generates substantial hydrostatic pressure in the gut, which, if unregulated, can draw water from the hemolymph, leading to dehydration and death^23^. Thus, successful feeding requires tightly coordinated decisions about intake, nutrient absorption, and excretion of sugar rich honeydew. In aphids, these decisions are influenced by the carbon-to-nitrogen ratio, the balance of essential to non-essential amino acids, and absolute sugar and amino acid levels^33–36^, although the underlying molecular mechanisms remain unknown.

To assess digested diet composition and regulate feeding behaviors such as intake and excretion, insects require dedicated chemosensory mechanisms. In phloem feeders, however, the nature of these mechanisms remains largely unknown. In *Drosophila melanogaster*, the gustatory receptor GR43a is expressed in the gut, brain, and proventricular ganglion neurons innervating the brain, foregut, and midgut. It is thought to monitor fructose levels in the diet and hemolymph, thereby regulating food intake, gut transit, and the release of neuropeptides or hormones involved in feeding behavior regulaion^37,38^. Homologs of *DmGR43a* in lepidopterans are also expressed in the digestive system and are proposed to serve similar functions^37,39,40^. In the beetle *Tribolium castaneum*, the mannitol and sorbitol receptor *TcGR20* enhances dietary intake in the presence of these sugars^41^. A key missing component, particularly given the role of amino acids in regulating phloem-feeding, is the mechanism of amino acids detection. To date, this has been characterized only in *D. melanogaster* and *Apis mellifera*, where members of the ionotropic receptor (IR) and GR families mediate attraction or aversion to dietary amino-acids^42–46^. Notably, in *D. melanogaster*, GRs that promote attraction to amino acids (GR5a, GR61a, GR64a) are also canonical sweet taste receptors responsive to specific sugars ^45^.

Here, we use the whitefly *B. tabaci* as a model to investigate how sugar and amino acid sensing contributes to the regulation of feeding behavior and diet intake in phloem feeders. We focus on BtabGR1, the only functionally characterized sweet GR in this species. Our results show that BtabGR1 integrates three environmental chemical cues: sucrose concentration, the presence of the essential amino acid arginine, and the pH values of the feeding solution. We discuss these findings in the context of the insect’s feeding environment and propose a molecular model for diet ingestion control based on an unusually high level of sensory integration for a single insect gustatory receptor.

## Results

### Arginine activates the sucrose receptor BtabGR1 under alkaline conditions

Building on findings in *D. melanogaster* showing that sweet taste receptors also mediate responses to amino acids^45^, we tested whether amino acids act as agonists of BtabGR1. We expressed *BtabGr1* in *Xenopus* oocytes and recorded receptor activity using the two-electrode voltage clamp system (Figure 1A). All 20 amino acids were applied (at 100mM or the highest soluble concentration) either individually or in chemically mixtures (hydrophobic, hydrophilic, acidic, or basic) to minimize potential interactions or neutralizing effects.

**Figure 1.**
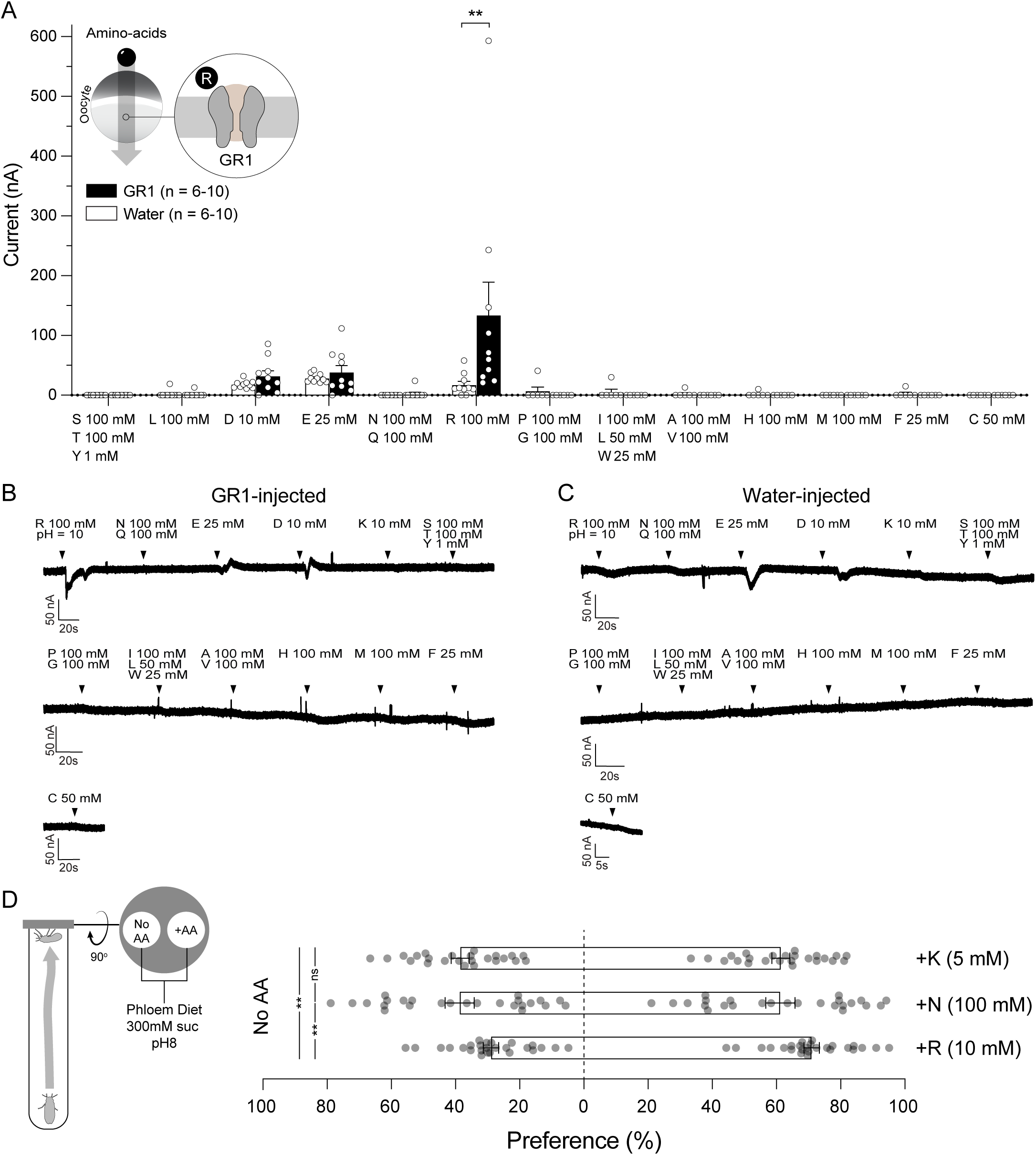
Arginine selectively activates BtabGR1 and enhances diet preference in whiteflies. **(A)** Mean current responses (±SEM, n = 6-10, one-way ANOVA, ∗∗*P* ≤ 0. 01) of *BtabGR1*-expressing and water-injected *Xenopus* oocytes to individual amino acids or amino acid mixtures. Amino acids were tested at 100 mM or at the highest soluble concentration. Representative current traces recorded from BtabGR1-expressing (**B**) or water-injected control (**C**) oocytes following exposure to individual amino acids or amino acid mixtures. Black arrows indicate solution application. **(D)** Percentage of adult whiteflies preferring amino acid-containing diets (lysine, K; asparagine, N; arginine, R) over amino acid-free diets (no amino acid, No AA) in dual-choice assays (mean ± SEM; n > 30 per treatment; \*\**P* ≤ 0.01). All diets contained 300 mM sucrose.

Among all tested samples, only arginine elicited a significant current response (**Figures 1A and 1B**) compared to water-injected controls (**Figure 1C**). Because arginine is a basic amino acid (solution pH 10), we next examined whether pH alone could account for receptor activation. Exposing oocytes to a pH range of 4 -10 did not produce significant responses (**Figure S1**), indicating that BtabGR1 does not function as a pH sensor and that alkaline conditions alone are insufficient for activation. Consistent with this, the other basic amino acids lysine (pH 9.5) and histidine (pH 7.5) did not elicit responses in either BtabGR1-expressing or control oocytes.

### Arginine enhances diet preference in whiteflies

To assess whether arginine sensing influences diet choice, we performed dual-choice bioassays with adult whiteflies. To approximate natural conditions, both options contained sucrose at phloem-like levels (300 mM, pH 8). One feeding site was supplemented with a single amino acid at physiologically relevant concentrations (phloem concentrations), while the other contained sucrose alone (**Figure 1D**). We compared responses to arginine with those to lysine (another positively charged amino acid) and asparagine, the most abundant amino acid in the phloem-sap of Fabaceae hosts of *B. tabaci* ^47^. Whiteflies showed a significant preference for diets containing lysine (5 mM; 61.36%, *P* < 0.0001) and asparagine (100mM; 61.199%, *P* = 0.0051) over sucrose alone. However, the preference for arginine (10 mM) was markedly stronger (70.95%, *P* < 0.0001) and significantly higher than for lysine or asparagine (*P* = 0.0035 and *P* = 0.0017, respectively).

### Arginine activation of BtabGR1 is pH dependent

To further characterize the interaction between BtabGR1 and arginine and assess the influence of pH on ligand potency, we performed concentration-response assays across four pH conditions (**Figure 2A**). pH 10 corresponded to the conditions used in the initial screening. pH 8 reflects the reported average of plant phloem sap^13–16^, pH 5.3 represents the reported average of the leaf apoplast^17–22^, and pH 6 approximates the gut environment of *B. tabaci*, as determined using bromothymol blue^48^ (**Figure S2**).

**Figure 2.**
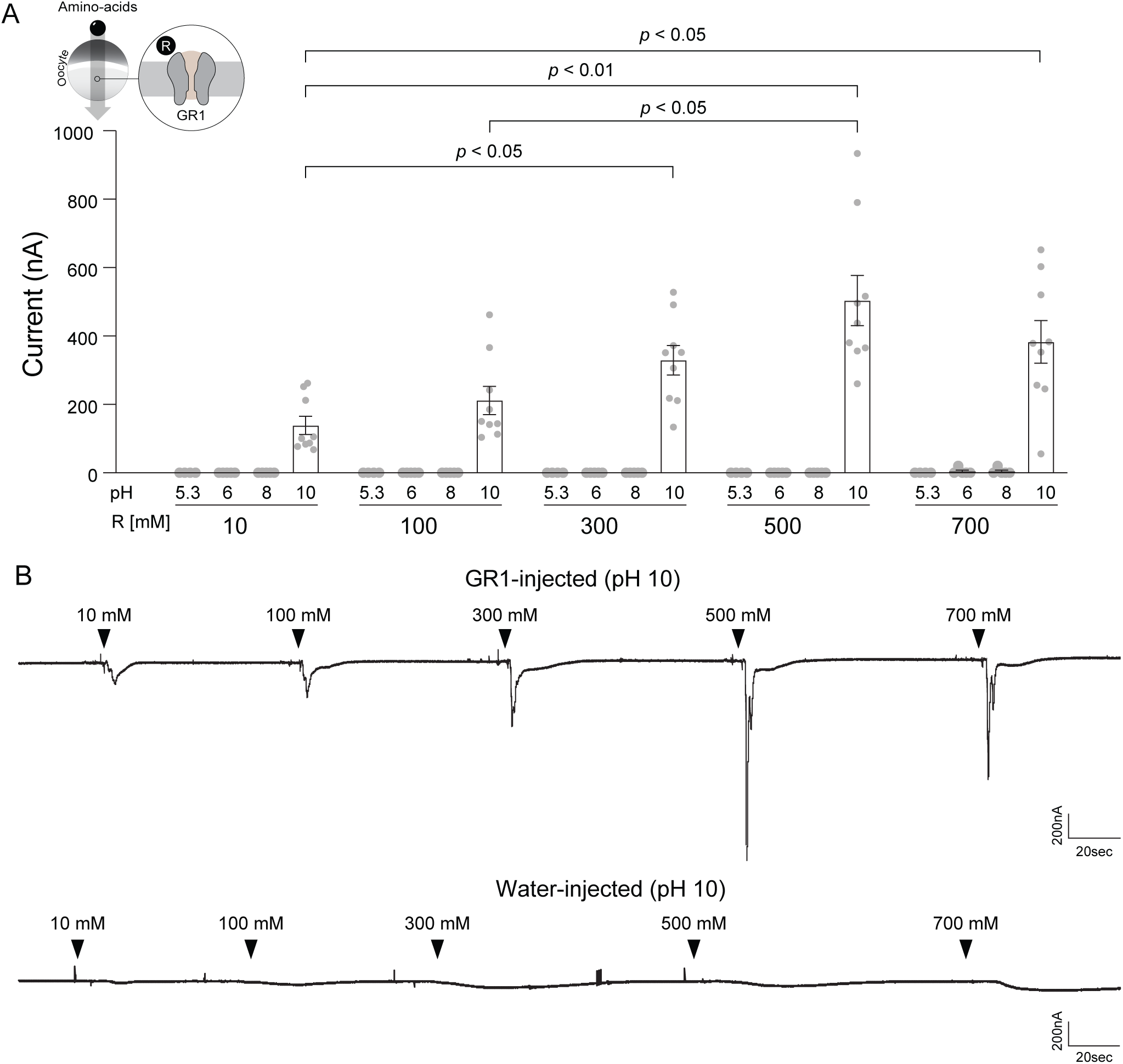
Arginine activation of BtabGR1 is pH-dependent. **(A)** Mean current responses (± S.E.M.) of BtabGR1-expressing *Xenopus* oocytes to arginine (10-700 mM) at pH 5.3, 6, 8 and 10 (n = 10 for pH 10; n = 4 for all other pH conditions; *t*-tests followed by FDR correction). **(B)** Representative current traces from BtabGR1-expressing oocytes in response to the indicated arginine concentrations at pH 10. Black arrows indicate solution application. All BtabGR1-expressing oocytes responded to the positive control (100 mM sucrose, pH 7.5; data not shown).

At pH 10, BtabGR1 exhibited a concentration-dependent response to arginine (**Figures 2A and 2B**). The current at 300 mM was significantly higher than that at 10 mM (*P* ≤ 0.05), with additional significant differences observed between 500 mM and 10 mM (*P* ≤ 0.01), 500 mM and 100 mM (*P* ≤ 0.01), and 700 mM and 10 mM (*P* ≤ 0.05). However, fitting the data to a Hill equation yielded poor fit (R^2^ square = 0.28 with fixed slope; R ^2^ = 0.3 with variable slope), indicating weak or non-classical ligand behavior and suggesting that arginine does not act as a high-affinity agonist under these conditions. In contrast, at physiologically relevant pH values (5.3, 6, and 8), arginine alone did not elicit detectable responses from BtabGR1 (**Figure 2A**). This strong pH dependence, together with the lack of activity under biologically relevant conditions, was unexpected and suggested that additional factors are required for arginine-mediated receptor activation.

### Selective enhancement of sucrose sensing by arginine at gut pH

To further resolve the pH-dependent interaction between arginine and BtabGR1, we examined how pH influences receptor responses to its primary ligand, sucrose, in the presence or absence of arginine (+R or -R). We hypothesized that this interaction would be most informative under conditions mimicking the environments encountered during feeding. Accordingly, we generated sucrose concentration-response curves (1-1,300 mM) at three physiologically relevant pH values: (i) apoplast (pH 5.3), (ii) phloem (pH 8), and (iii) gut (pH 6). Arginine was added at biologically relevant concentrations (10 mM for phloem and gut; 0.3 mM for apoplast; see Introduction).

At apoplast pH (pH 5.3), neither EC_50_ values (-R: 469.5 mM, +R: 206.9 mM, *P* = 0.28) nor response amplitudes differed significantly between treatments, including the maximal response (**Figure 3A**). At phloem pH (pH 8), the -R curve was slightly left-shifted and reached higher amplitudes than the +R curve (**Figure 3B**). However, EC_50_ values did not differ significantly (-R: 348.7 mM, +R: 410.2 mM, *P* = 0.51), and response amplitudes differed at only two sucrose concentrations (100 and 300 mM) (**Figure 3A**). The maximal response at 1,300 mM sucrose was higher in the -R condition than the +R condition (- R:11,627 nA, +R: 8973 nA), but the difference was not significant (*P* = 0.0586). Overall, these data indicate a modest reduction in sucrose responsiveness in the presence of arginine at pH 8.

**Figure 3.**
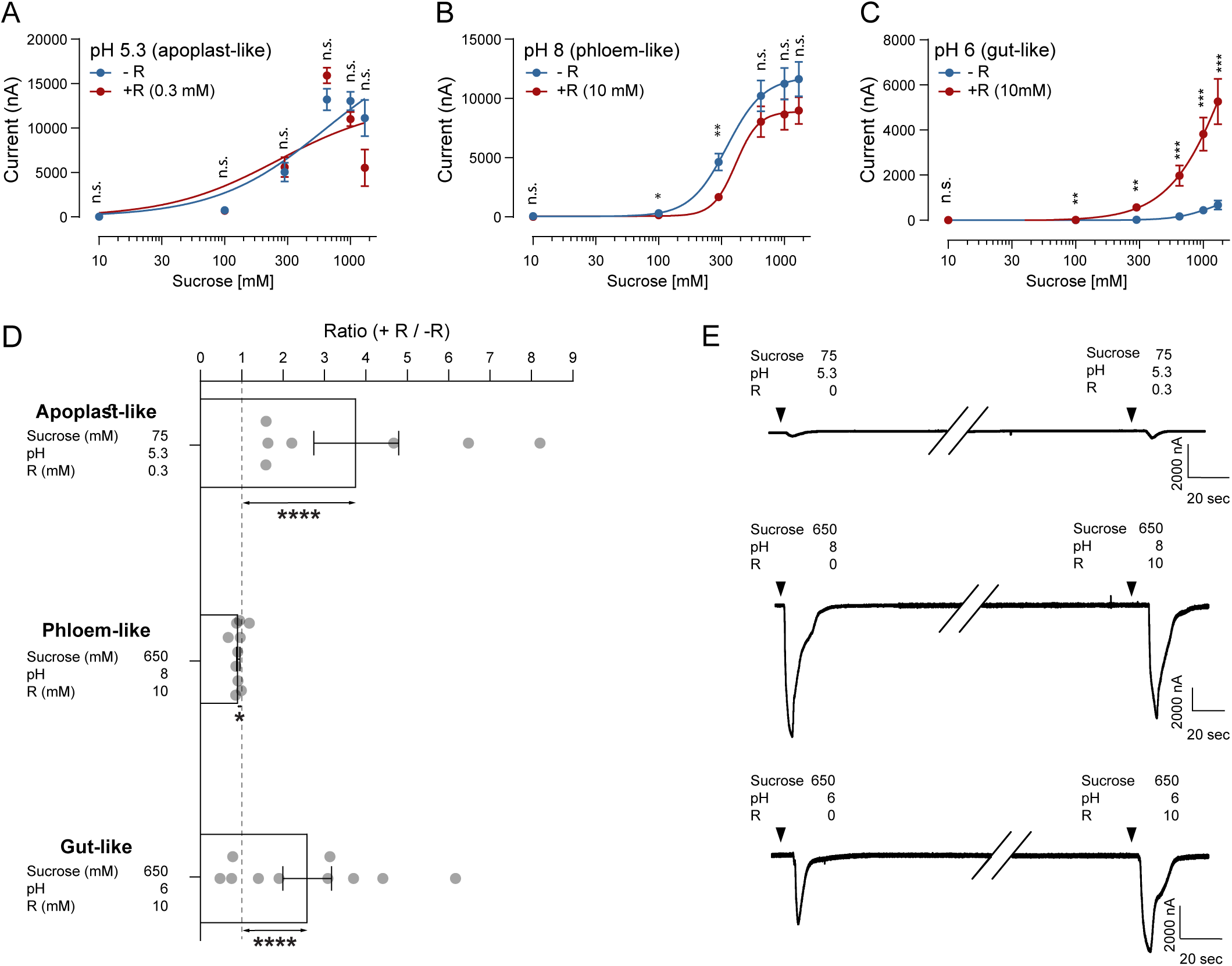
Arginine enhances BtabGR1-mediated sucrose sensing in acidic environments. **(A-C)** Sucrose concentration-response curves (CRCs: 10, 100, 300, 650, 1000, 1300 mM) recorded in the presence (+R) or absence (-R) of arginine at different apoplast-, phloem- and gut-like pH conditions (mean response ± S.E.M, one-way ANOVA; *** *P* < 0.001, ** *P* < 0.01, \**P* < 0.05, ns = non-significant). **(D)** Pairwise comparisons of current responses to apoplast-, phloem- and gut-like solutions in the presence or absence of arginine (+R and -R, respectively). Bars represent the mean -R /+R response ratio ± S.E.M. (n = 7-10). Ratios were tested against a theoretical value of 1 using *t*-tests. **** *P* < 0.0001, \**P* < 0.05. **(E)** Representative current traces from BtabGR1 expressing *Xenopus* oocytes exposed to gut-, apoplast- and phloem-like solutions with or without arginine. (Full traces including responses to 100 mM sucrose which were recorded between treatments for normalization across oocytes are available in Figure S3). Black arrows indicate solution application.

In contrast, a striking effect emerged under gut-like conditions (pH 6). Here, Hill fits were poor (+R, R^2^ = 0.5; -R, R^2^ square = 0.6), precluding reliable EC_50_ estimation (**Figure 3C**). Nevertheless, the -R curve was markedly right-shifted and flattened, with significantly lower responses at all sucrose concentrations above 10 mM. These data suggest that sucrose alone is a weak agonist at gut pH. In contrast, responses observed in the presence of arginine were consistently elevated, indicating that arginine acts as a pH-dependent positive modulator that potentiates sucrose-evoked activation under gut conditions.

To further assess physiological relevance, we performed pairwise comparisons using solutions mimicking the apoplast (pH 5.3, 75 mM sucrose), phloem (pH 8, 650 mM sucrose), and gut (pH 6, 650 mM sucrose), each tested with or without arginine (**Figure 3D and 3E**). Under apoplast-like conditions, arginine (0.3 mM) enhanced receptor responses nearly fourfold (+R / -R = 3.767, *P* < 0.0001), indicating enhanced sucrose detection. Under phloem-like conditions, arginine (10 mM) caused a small but significant reduction in response (+R / -R = 0.914., *P =* 0.0333). A significant effect was observed again under gut-like conditions, where arginine (10 mM) more than doubled sucrose-evoked currents (+R / -R = 2.58, *P* < 0.0001). Together, these results support the hypothesis that arginine acts as a context-dependent modulator of BtabGR1, selectively enhancing sucrose sensing under acidic apoplast- and gut-like conditions.

### Role of BtabGr1 and arginine in diet ingestion

In a previous study, we showed that *BtabGr1* is expressed in body regions associated with sucrose sensing in phloem feeders, namely the head and abdomen^28^. RNA-seq analysis conducted in this study further revealed that *BtabGR1* is expressed in gut and mouthpart tissues (**Figure S4**). The strong modulatory effect of arginine on sucrose responses at gut pH suggested that arginine in the gut lumen may contribute to the regulation of diet ingestion. To test this hypothesis, we developed an assay using honeydew excretion as a proxy for diet intake (**Figure 4A**). Adult whiteflies were first fed for 72h on a pH 8 solution containing 300 mM sucrose and 0.5 µg/µL dsRNA targeting either *BtabGR1* (*dsBtabGr1*) or *GFP* (*dsGFP*) as a control. They were then transferred to solutions containing 300 mM sucrose and the same dsRNA, with or without 10 mM arginine. This design yielded four treatments differing in arginine supplementation and dsRNA identity. qRT-PCR confirmed effective gene silencing (**Figure 4B**). In the absence of arginine, *BtabGr1* transcript levels were reduced by 38.2% in *dsBtabGr1*-fed insects compared to *dsGFP* controls (*P* = 0.04). In presence of arginine, transcript levels decreased by 31%, although this reduction was not statistically significant (*P* = 0.128). Analysis of honeydew excretion revealed significant effects of both arginine (*P* < 0.0001) and dsRNA (*P* = 0.047) main treatments. Pairwise comparisons indicated that silencing of *BtabGr1* in the absence of arginine (-R) significantly enhances diet intake (*P* = 0.017). On the other hand, in the presence of arginine (+R), there was no significant difference in diet intake between *dsBtabGr1*- and *dsGFP-*fed insects (*P* = 0.647), suggesting some mild-interaction between the silencing of *BtabGr1* and arginine presence/absence. (**Figure 4C**).

**Figure 4.**
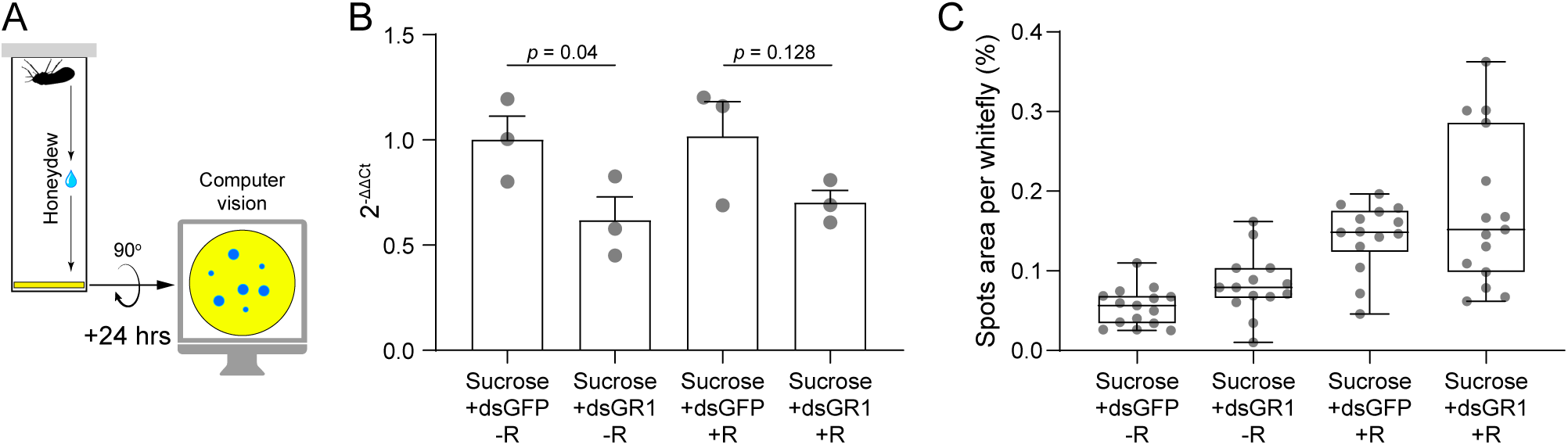
Arginine and *BtabGr1* regulate diet ingestion in whiteflies. **(A)** Schematic of the diet intake assay. The assay consisted of a feeding site positioned in the lid of a tube and a SpotOn paper disc placed at the bottom. Honeydew droplets falling onto the paper produced blue spots, the relative area of which was used as a measure of honeydew excretion, and consequently, a proxy for diet ingestion. **(B)** qRT-PCR analysis of *BtabGR1* expression in whole body homogenates of adults fed diets containing dsRNA targeting *BtabGr1* (*dsGr1*) or *Gfp* control dsRNA (*dsGfp*) (mean ± SEM, n = 3). **(C)** Ratio of honeydew-spotted area to total disc area (mean ± S.E.M, n = 14 -15) in adults fed 300 mM sucrose diets containing *dsGr1* or *dsGfp*, with or without 10 mM arginine (+R or -R, respectively). For both **(B)** and **(C)**, statistical significance was determined using two-way ANOVA followed by pairwise comparisons with FDR correction.

## Discussion

In this study, we expand our understanding of the role sweet GRs play in phloem-feeding using the whitefly *B. tabaci* as a model. In a previous study, we showed that BtabGR1 is finely tuned to sucrose detection and plays a key role in feeding-site allocation, which was demonstrated by artificial diet choice assays and gene silencing experiments^28^. Here, we extend this framework by examining whether BtabGR1 also detects additional ligands, contributes to the assessment of feeding-site quality, and regulates diet-intake.

Our results reveal an unexpectedly complex interaction between BtabGR1 and three environmental factors: sucrose concentrations, the presence of the essential amino acid arginine and pH values of the feeding solutions. Notably, arginine emerges as a pH-dependent modulator of sucrose sensing. We interpret these findings from two complementary perspectives. First, we propose that integration of these extracellular cues influences the evaluation of chemical signals encountered during probing, contributing to a directional stylet movement towards optimal feeding sites. Second, we suggest that arginine levels in the gut lumen provide feedback on diet quality, thereby contributing to the regulation of ingestion.

Although sucrose concentration-response curves showed no overall differences between -R and +R treatments at phloem-like (pH 8) or apoplast-like (pH 5.3) conditions, analysis at biologically relevant sucrose concentrations revealed more nuanced effects. At pH 8, the addition of 10 mM arginine led to a modest (∼9%) but significant reduction in BtabGR1 response. While the functional significance of this effect remains unclear, it may reflect more physiologically realistic receptor behavior under natural conditions where the amino acid is present, compared with artificial diets lacking arginine entirely^36^. Stronger effects were observed under apoplast-like conditions. At pH 5.3, the presence of 0.3 mM arginine increased receptor responses to sucrose nearly fourfold, indicating a functional role for arginine in the apoplast. Like other amino acids, arginine is loaded into the phloem in most crop plants via apoplastic transport, which can result in higher amino acid concentrations at the extracellular space surrounding the sieve element-companion cell complex^7–9^. Although direct measurements are lacking, apoplastic arginine concentrations are expected to reach up to ∼0.5 mM^12^. Combined with our previous findings that BtabGR1 enables tracking of sucrose gradients^28^, these data suggest that simultaneous sensing of sucrose and arginine may enhance navigation toward the feeding site.

Taking together the concentration-response curves and pairwise comparison results, the most pronounced effect of arginine on BtabGR1 activity was observed under gut-like conditions (pH 6). At this pH, sucrose alone acted as a weak agonist, with robust receptor activation largely depending on the presence of arginine. Similar mixture-dependent effects have been reported in nectar-feeding hummingbirds, where certain amino acids show little activity alone but enhance receptor responses when combined with nectar sugars^49^. Consistent with the robust response of BtabGR1, our diet intake bioassays demonstrated that the presence of arginine in the gut lumen significantly enhanced ingestion. In *D. melanogaster*, gut sugar levels are monitored by neurons expressing sugar receptors that project into the lumen^37^, as well as by enteroendocrine cells expressing receptors such as Gr64a (sucrose/maltose) and Gr43a (fructose)^50^. Activation of these receptors triggers the release of neuropeptide F, which acts on the corpora cardiaca and insulin-producing cells to regulate feeding and sugar satiety^51^. Given that BtabGR1 is expressed in the gut of *B. tabaci*, it may function in a similar neuroendocrine framework, with the exception of ability to integrate signals from both sucrose and arginine into a single output. Such integration offers a potential mechanistic explanation for earlier observations in phloem-feeders, showing that nitrogen limitation drives increased food intake in aphids, and that this compensatory response depends not simply on total amino acids levels, but on the detection of specific amino acids or their combinations^36^.

An important question arising from these findings is why arginine, in particular, would be sensed to regulate diet ingestion in *B. tabaci*. One possibility is that additional amino acids are also detected by GRs that have not been characterized yet, meaning that arginine may not be unique. Nevertheless, a specific capacity to detect arginine is biologically plausible. Arginine is chemically distinct in its guanidinium group which remains strongly protonated across the pH range relevant to whitefly feeding (pH 5.3-8)^52,53^. This stable cationic character may promote consistent interactions with receptor binding sites, across changing pH conditions. as shown for L-arginine-specific taste receptors in catfish^54^. Further support for this view comes from studies in other insects showing that arginine is frequently detected by amino acid-responsive receptors. For example, in *D. melanogaster*, GR5a, GR61a, and GR64f, as well as GR10 in *A. mellifera*, respond to arginine alongside other amino acids. Similarly, several ionotropic receptors involved in amino acid sensing in *D. melanogaster* (IR25a, IR51b, IR76, and IR60c) detect arginine^42,43,45,46,55^. Together, these findings suggest that arginine is a prominent cue in insect gustatory systems.

Arginine is also nutritionally informative. As an essential amino acids, it is relatively scarce in phloem-sap, comprising ∼3% of the total amino acid pool, with essential amino acids collectively accounting for only 21%^47^. Phloem-feeding insects compensate for this imbalance through obligate symbionts capable of synthesizing both essential and non-essential amino acids^56^. In the aphid *Acyrthosiphon pisum*, symbiont-mediated arginine synthesis is suppressed when dietary arginine is sufficient^36^. Arginine also regulates nitrogen metabolism by inhibiting the glutamine transporter ApGLNT1, which controls glutamine uptake into bacteriocytes^57^. Because glutamine serves as a precursor for the synthesis of the remaining 19 amino acids, arginine levels may act as an indicator of overall amino acid availability in the hemolymph and gut lumen^57^. Consistent with this view, analysis across different legume hosts, demonstrated that phloem arginine concentrations show the strongest positive correlation with aphid performance ^11^. Finally, arginine has the highest nitrogen-to-carbon ratio among amino acids, making it an especially valuable nitrogen source^58^. Together, these characteristics identify arginine as a metabolically and nutritionally strategic molecule, well suited to function as a signal of dietary quality in phloem-feeding insects.

Collectively, these findings support an expanded model for the role BtabGR1 plays in whitefly feeding behavior, extending our 2023 study^28^ (**Figure 5**). In the apoplast, sucrose gradients are thought to guide stylet progression toward the phloem^28^. Our data suggest that a parallel arginine gradient, potentially shaped by similar transport mechanisms (apoplastic loading followed by active uptake into the sieve-elements)^2,8,9^, may further enhance receptor activation as the stylet advances. Integration of sucrose and arginine signals could thus improve the precision and efficiency of feeding-site localization. At phloem pH, BtabGR1 shows maximal sensitivity to sucrose, enabling reliable recognition of sieve-elements following stylet penetration. Once ingestion begins, BtabGR1 may also contribute to diet evaluation within the gut. Under gut-like pH conditions, receptor activity is strongly enhanced by arginine, which acts as a positive modulator. This interaction may promote increased consumption when both sucrose and arginine are present. In this respect, it is important to note that our silencing assays adds an additional and not completely resolved level of complexity to the expanded model. In the absence of arginine, *BtabGr1* silencing leads to enhanced diet intake, likely due to a compensatory mechanism in which lower receptor activity is interpreted as lower sucrose sensing (i.e. lower sucrose concentrations), leading to higher consumption^35^. In the presence of arginine, gene silencing initiates the activity of two contradicting forces. Again, lower receptor activity is interpreted as lower sucrose sensing leading to higher consumption, but at the same time, the enhancing effect of the presence of arginine is compromised in *dsBtabGr1-*fed insects. This leads to equal diet intake by silenced and non-silenced insects. Further experiments are required to resolve the delicate relationship between the two opposing effects and the contribution of their balance to feeding-site acceptance. In any case, it can be stated that the combined effects of sucrose, arginine, and pH on BtabGr1 define a highly interdependent sensory system, representing the most complex example to date of environmental signal integration by a single insect chemoreceptor.

**Figure 5.**
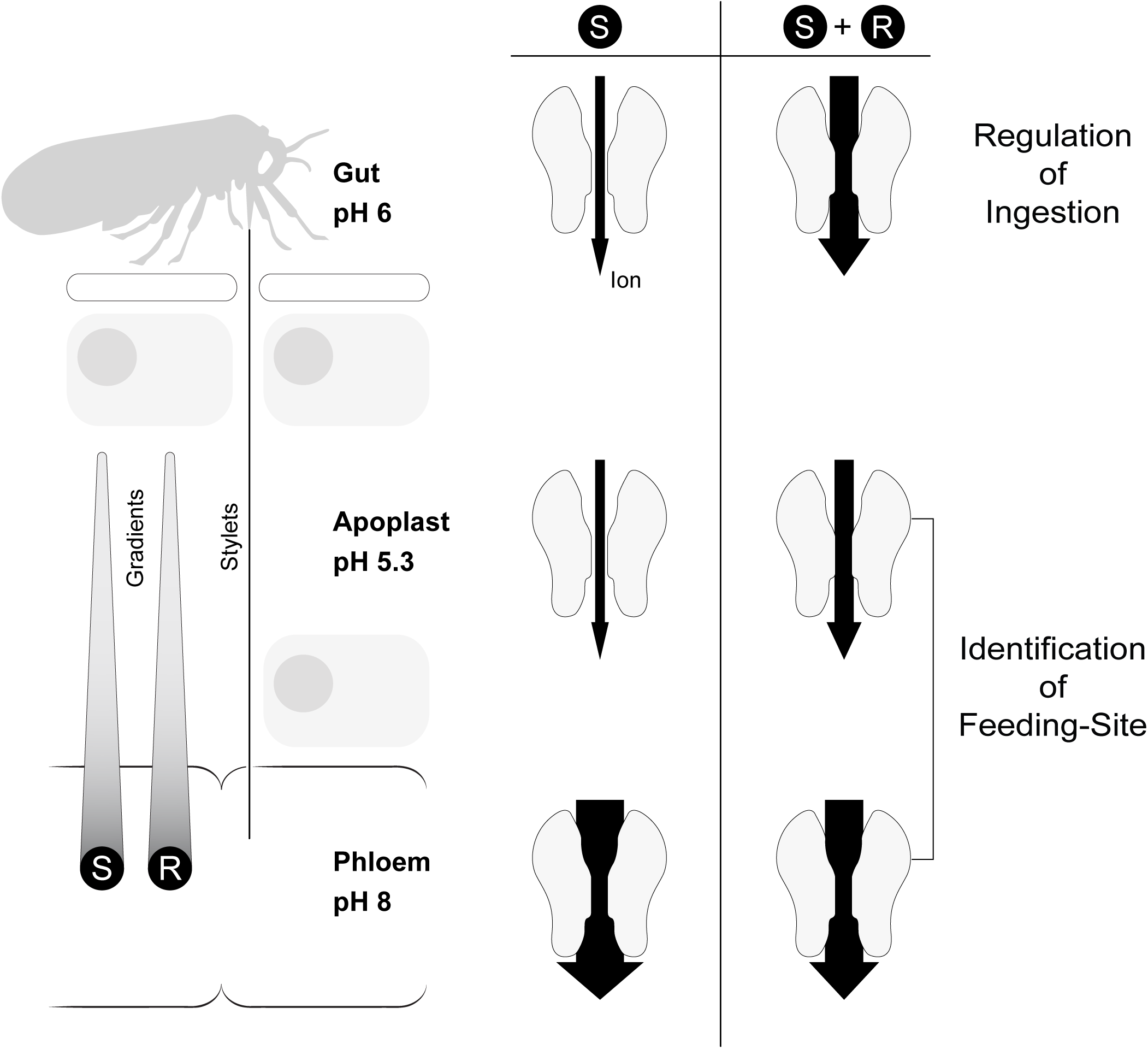
A model for pH-dependent integration of feeding cues by BtabGR1. In this model, arginine acts as a pH-dependent modulator of BtabGR1, selectively enhancing sucrose sensing (indicated by increased thickness of black arrows) under apoplast- and gut-like conditions. **(Bottom)** In the phloem, BtabGR1 exhibits maximal sensitivity to sucrose, largely independent of arginine. **(Middle)** In the apoplast, elevated sucrose and arginine concentrations in the extracellular space surrounding the sieve element–companion cell complex enhance receptor activation and may improve the efficiency of feeding-site localization. **(Top)** Under gut-like conditions, arginine strongly potentiates receptor activity, enhancing sucrose sensing and potentially promoting increased diet ingestion when both compounds are present.

## Materials and methods

### *Bemisia tabaci* population and rearing

A population of the local (Israeli) species of the whitefly *B. tabaci* (the MEAM1 species) was used. The population was established in 2019 by collecting ∼ 6,000 adults from four tomato and watermelon open fields. Since its collection, the population has been maintained on both host plants (six cages contributing equally to each new generation) in the laboratory insectaries (28 ± 2 ℃ and a 14:10-hr light:dark cycle). All described bioassays were conducted on a sub-population of the one described above reared for more than 20 generations only on cotton plants.

### Determining gut pH using bromothymol staining

Newly emerged (∼3 days-old) adults were collected and immediately frozen in liquid nitrogen. Each individual was then dissected in 5 µl of 0.1 x Acetate Buffer Solution (ABS; Sigma-Aldrich), separating the gut sample from the rest of the body. Next, the gut sample was moved into bromothymol blue solution in ABS. After the color of the gut stabilized, a photo was taken under a binuclear. The gut color in the photo was compared to a standard scale of bromothymol blue at changing pH^48^.

### Dual choice assays

A laboratory custom-designed dual choice-chamber system (30 x 200 mm glass vials) was used. Diet solutions were placed between two layers of Parafilm (4 x 4 cm) stretched over a 3D-printed lid with two open round windows. It was assumed that the Parafilm is perceived by the insects as the plant cuticle ^32^. Moreover, the subsequent steps in which stylet penetration of the Parafilm, probing, and sampling take place, were considered as a mimic of apoplast and/or sieve-element evaluation. Therefore, a decision to leave the test solution or to continue ingesting was interpreted as rejection or acceptance of specific nutritional conditions in plants.

Forty newly emerged (∼3 days-old) couples (40 males and 40 females) were collected into a glass vial using a keyboard vacuum cleaner. Next, the dual choice lids containing the diet solutions were placed on top of the vial and sealed together using a parafilm strip. Each vial was covered with aluminum foil, leaving only the lid exposed to light. After 24 h (at 28 ℃ and 14:10 L:D photoperiod), the number of settled adults in each of the two feeding sites was recorded. To control for potential position effect, the lids of the replicates were rotated by 180° so that the position of the two feeding sites was alternated between replicates.

### Statistics applied to dual choice assay

Differences in diet preference (between the diets offered in the two feeding sites of the chambers) were tested separately for each choice assay using a paired *t*-test and a null hypothesis of equal distribution (50% of the individuals feeding at each site). Statistical significance was assumed at *P* ≤ 0.05. When more than one dual choice assay was done, and a comparison between the preferences in each test was needed, the significance of the differences in diet preference was analyzed using a one-way ANOVA model followed by a-priori pairwise comparisons and false discovery rate (FDR) correction (R v. 4.5.2). Statistical significance was assumed at *P* ≤ 0.05. All described statistical analyses were conducted using JMP Pro 18.0 (SAS Institute, Cary, NC).

### Xenopus laevis rearing

The *Xenopus laevis* females (pigmented *Xenopus laevis*) were obtained from Nasco (Fort Atkinson, WI, USA). The frogs were reared in a XenoPlus amphibia housing system (Tenciplast, West Chester, PA, USA) under controlled water conditions (pH = 7.6, Temp = 18.2 °C, conductivity = 1100 µS). The frogs were fed three times a week with adult *Xenopus* irradiated diet (Zeigler, Gardners, PA, USA, Prod. No. 316518-18-2412).

### *BtabGR1* cloning and mRNA expression in *Xenopus laevis* oocytes

*BtabGR1* cDNA was cloned into pSP64T (Addgene, CAT# 15030, Watertown, MA, USA), using Takara-Clontech infusion kit (CAT# 638909). *In-vitro* transcription was done using the mMESSAGE Mmachine sp6 Transcription kit (CAT# AM1340, Thermo Fisher, Waltham, MA, USA). Stage IV-VI oocytes of *X. laevis* were harvested, mechanically disrupted by hand, and treated with a 10 mg/ml collagenase solution (Sigma Aldrich, CAS) at 18 ℃ for 25 minutes at 60 RPM. Following collagenase treatment, oocytes were washed 5 times in ND96 solution (96 mM NaCl, 2 mM KCl, 5 mM MgCl2, and 5 mM HEPES, 1L double-distilled water, pH 7.6). Subsequently, oocytes were rinsed in a washing solution (ND96 solution supplemented with 100 μg/mL gentamicin). Finally, oocytes were washed 5 times with incubation medium (ND96 solution supplemented with 0.8 mM CaCl2, 5% dialyzed horse serum, 50 μg/mL tetracycline, 100 μg/mL streptomycin and 550 μg/mL sodium pyruvate). Oocytes were allowed to recover overnight prior to injection with 27.6 nL cRNAs (3 μg/μl), followed by incubation at 18 ℃ for 3 days in 24 well-plates containing 1 ml incubation medium.

### Pharmacology -experimental system

Whole-cell currents were monitored and recorded using the two-electrode voltage clamp (TEVC) technique. Holding potential was maintained at -80 mV using an OC725C oocyte clamp (Warner Instruments, LLC, Hamden, CT, USA). Oocytes were placed in a RC-3Z oocyte recording chamber (Warner Instruments, LLC, Hamden, CT, USA) and exposed to four-second-long stimuli. All compounds were solubilized in ND96 buffer supplemented with 0.8 mM CaCl2. Data acquisition was carried out with Digidata 1550A and pCLAMP10 (Molecular Devices, Sunnyvale, CA, USA).

### Pharmacology -ligand screening

To assess the amino acid selectivity of BtabGR1, 14 amino acids mixes were used as presented in **Figures 1A to 1C**. Currents were allowed to return to baseline between ligand applications. The screening protocol included administration of the mixes in forward and reverse orders to control for position effects. The mean current responses (±SEM, n = 6-10) of *BtabGR1*-injected oocytes were compared to those of water-injected oocytes (one-way ANOVA).

### Pharmacology – concentration response curves (CRCs)

Seven solutions of either rising arginine or sucrose concentrations were administered for four seconds each, and whole cell currents were allowed to return to baseline. The resulting concentration-response curves, EC50 interpolation, EC comparison, and statistical analyses were conducted using GraphPad Prism 8 (GraphPad Software Inc., La Jolla, CA, USA). One-way ANOVA was used to compare response values between two conditions at the same concentration (+R vs. -R), and multiple *t*-tests were used to compare response values between different arginine concentration and maximal recorded response (Emax). Comparisons were conducted using JMP Pro 18.0 (SAS Institute, Cary, NC). FDR corrections were conducted using R v. 4.5.2.

### Pharmacology – response comparison

Sucrose solutions in selected pH with (+R) and without (-R) arginine and 100mM sucrose only solutions in between, were administered for four seconds each, and whole cell currents were allowed to return to baseline. The response value ratio of -R/+R was calculated for each oocyte, and the mean ratio was tested to be different from 1 using *t*-tests. Analysis was conducted using JMP Pro 18.0 (SAS Institute, Cary, NC). FDR corrections were conducted using R v. 4.5.2.

### dsRNA design

The dsRNA constructed against a 370 nucleotide fragment of the green fluorescent protein (GFP) gene of the jellyfish *Aequorea victoria* was described in Luo et al. 2017^59^. For designing the dsRNA that targets *BtabGR1*, we first used the gene’s coding sequence as input for DSIR ^60^. Highly ranked siRNAs were aligned to the *BtabGR1* coding sequence, and a region (501 nt) containing high abundance of mapped siRNAs was identified. Next, all possible 21-mers of the selected region were extracted using Splitter (splitter-size 21 -overlap 20) ^61^. Usearch v.11 (usearch -search_oligodb insect -dbfile -strand both) ^62^ was used to verify that all 21-mers do not have an exact match or one mismatch to off-target genes in the MEAM1 genome (using the RefSeq annotated genome, GCF_001854935.1, in NCBI).

### Diet intake assays combined with gene silencing -experimental design

One hundred newly-emerged (∼3 days-old) couples (100 males and 100 females) were fed in no-choice glass vials on a solution containing 300 mM sucrose and 0.5 µg/µl of double stranded RNA (*dsBtabGR1* or *dsGFP* as control). After 72 hours, 16 individuals were transferred into smaller vials (18 x 50 mm glass vials) and left to feed for 24 h on a non-choice lid filled with solutions containing 300 mM sucrose, 0.5 µg/µl of double stranded RNA (*dsBtabGR1* or *dsGFP* as control) either with or without 10 mM of L-arginine. To quantify the amount of honeydew excreted (diet intake) during adult feeding on the different treatments, the bottom of each small vial was covered with a 16 mm diameter circular piece of SpotOn paper (CAT# 32920, innoquest, Inc. USA). The SpotOn paper is designed to spot moisture of any kind by switching color from yellow to blue in any part of the paper exposed to moisture or liquid. In our study, the honeydew excreted by the feeding whiteflies falls directly on the SpotOn paper, creating a blue dot sign on the paper.

### Quantification of honeydew excretion (diet intake)

Quantification of the blue area on the SpotOn papers involved few steps. First, a photo under a binuclear with a greyscale filter was taken. Next, the “Autocontext (2-stages)” workflow in ilastik v1.4.0 ^63^ was used for training an AI image analysis model to recognize and differentiate between the background of the photo, the paper disc, and the blue spots on it. The model was then used to process the photos and to create a black and white image where the spots are represented by black and the paper disc by white. The images served as an input to a transcript code (supplementary material – “spot-area-counter.ijm”) that run on Fiji software ^64^. The transcript calculates the total area of the disc and the area of spots on the disc. Then, for each disc, the ratio of spotted area to the whole disc area is calculated and normalized to the number of living whiteflies at the end of the assay. The effects of the dsRNA identity (*dsBtabGR1* or *dsGFP*) and the presence/absence of arginine in the diet on the ratio of the averaged spotted area were compared between treatments using a two-way ANOVA model. Analysis was conducted using JMP Pro 18.0 (SAS Institute, Cary, NC).

### Real-Time PCR for gene silencing quantification

For quantification of the silencing effect on *BtabGR1* expression in the diet intake assay, RNA was extracted from the survivors of the experiment, three samples per treatment (*dsGFP* -R, *dsGFP* +R, *dsBtabGR1* -R, *dsBtabGR1* +R), 30 individuals per sample, using the Isolate II RNA mini kit (Meridian Bioscience, Ohio). RNA quality and quantity were evaluated using a NanoDrop 2000 spectrophotometer (Thermo Fisher Scientific, Massachusetts). cDNA was synthesized using 500ng RNA from each sample and the Verso cDNA synthesis kit with Oligo-dT primer (Thermo Fisher Scientific, Massachusetts). The expression level of *BtabGR1* in *dsBtabGR1*-or *dsGFP*-fed adults was determined using the CFX Connect Real-Time PCR System (BIO-RAD, California). The *B. tabaci* endogenous ribosomal protein L13a (RPL13A) was used as a reference gene ^65^. The qRT-PCR conditions were adjusted for the amplification efficiencies of the target and endogenous genes to be in the log range of 1.9–2.1 and were optimally set to a master mix containing 5µl iTaq Universal SYBR Green Supermix (BIO-RAD, California), 0.5µl of both forward and reverse primers (primers 1-4, Table S2, 2pmol/µl), 2µl DDW, and 2µl cDNA template. qRT-PCR thermal conditions were: 95 ℃ for 2min, followed by 40 cycles of 95 ℃ for 5 sec and 60 ℃ for 30 sec, and an ending cycle of 95 ℃ for 5 sec, 65 ℃ for 5 sec, and 95 ℃ for 30 sec. Quantification of the *BtabGR1* gene expression levels was conducted following the ΔΔCt method (Applied Biosystems). A one-way ANOVA model followed by pairwise comparisons and FDR correction was used to determine the significance of the differences between the ΔCt means of treatment and control samples (*P* ≤ 0.05). Analysis was conducted using JMP Pro 18.0 (SAS Institute, Cary, NC).

### Gut and mouthparts RNA isolation and Illumina sequencing

Groups of 100 Newly emerged (1–7 days old) adult whiteflies from colonies reared on brussels sprouts were collected into 1.7 ml tubes and plunged into liquid nitrogen. This pool of tubes was deep frozen at -80 ℃ for future tissue collection. The whiteflies were dissected to get either isolated guts or mouthparts. Three samples were collected for each sex (females and males) and organ (gut and mouthparts). For each gut sample, 50 individuals were dissected, and 100 individuals were dissected for every mouthpart sample. All samples were homogenized with the first buffer (NucleoSpin RNA XS, Macherey-Nagel) using a dedicated drill. Total RNA was extracted from the homogenates according to the manufacturer’s instructions (NucleoSpin RNA XS, Macherey-Nagel). TruSeq RNA Library v2 libraries were constructed by Macrogen Inc. One sample of female guts did not yield a library, hence 11 libraries were sequenced with an Illumina sequencer platform. On average, approximately 40 million 151-bp pair-ended reads per sample (biological replicate) were produced.

### Gut and mouthparts RNA-seq processing and per-position read coverage profiling

All 11 gut and mouthpart total-RNA samples were mapped with STAR v2.7.11b^66^ to a curated reference of the annotated *B. tabaci* putative sweet-taste GRs (BtabGR1–4^28^, BtabGR5^67^). As the reference was not genome-wide, gene-level quantification was enabled (--quantMode GeneCounts), and raw counts were taken from the unstranded column of STAR’s ReadsPerGene.out.tab. Per-base coverage profiles were generated for each GR gene in every sample. Reads overlapping each gene were extracted with Rsamtools::ScanBamParam and converted to per-base depth with GenomicAlignments::coverage, then plotted with ggplot2::geom_area, faceted by condition (GF -guts of females, GM -guts of males, MF-mouthparts of females, MM -mouthparts of males.) with replicates left-to-right and a free y-axis per panel to compare coverage shape independent of library depth. Analyses used R v4.3.3 with the Bioconductor packages GenomicAlignments, Rsamtools and GenomicRanges.

