## Supplementary figures and images for "An Integrative Taste Receptor Links pH and Amino Acids to Sugar Sensing in *Bemisia tabaci*"

### Supplemental Figure 1

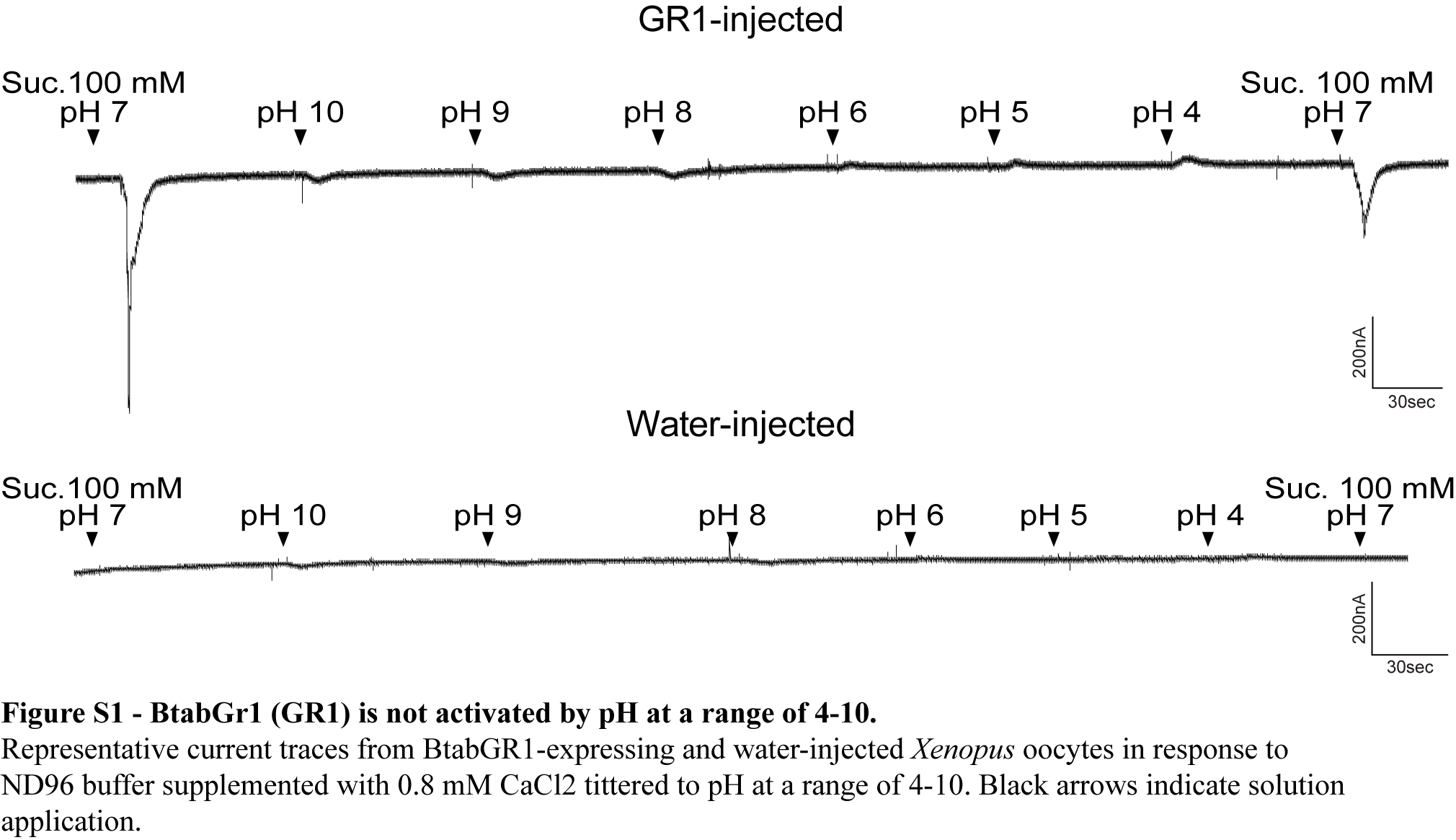

### Supplemental Figure 2

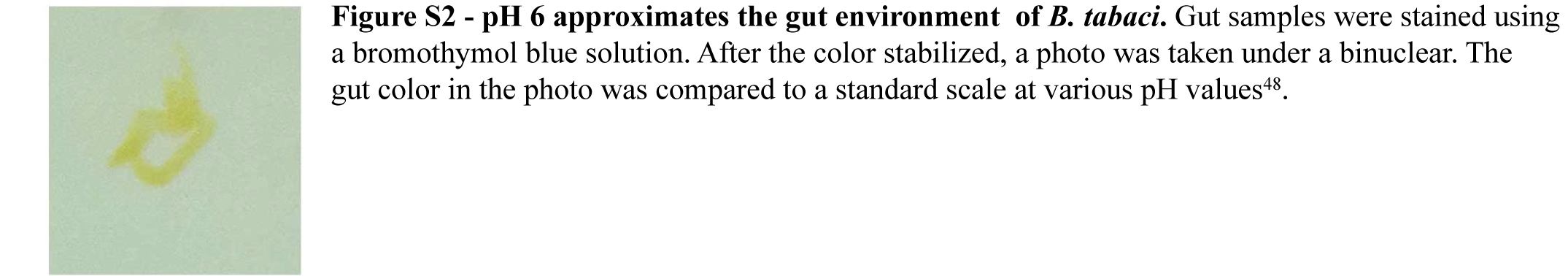

### Supplemental Figure 3

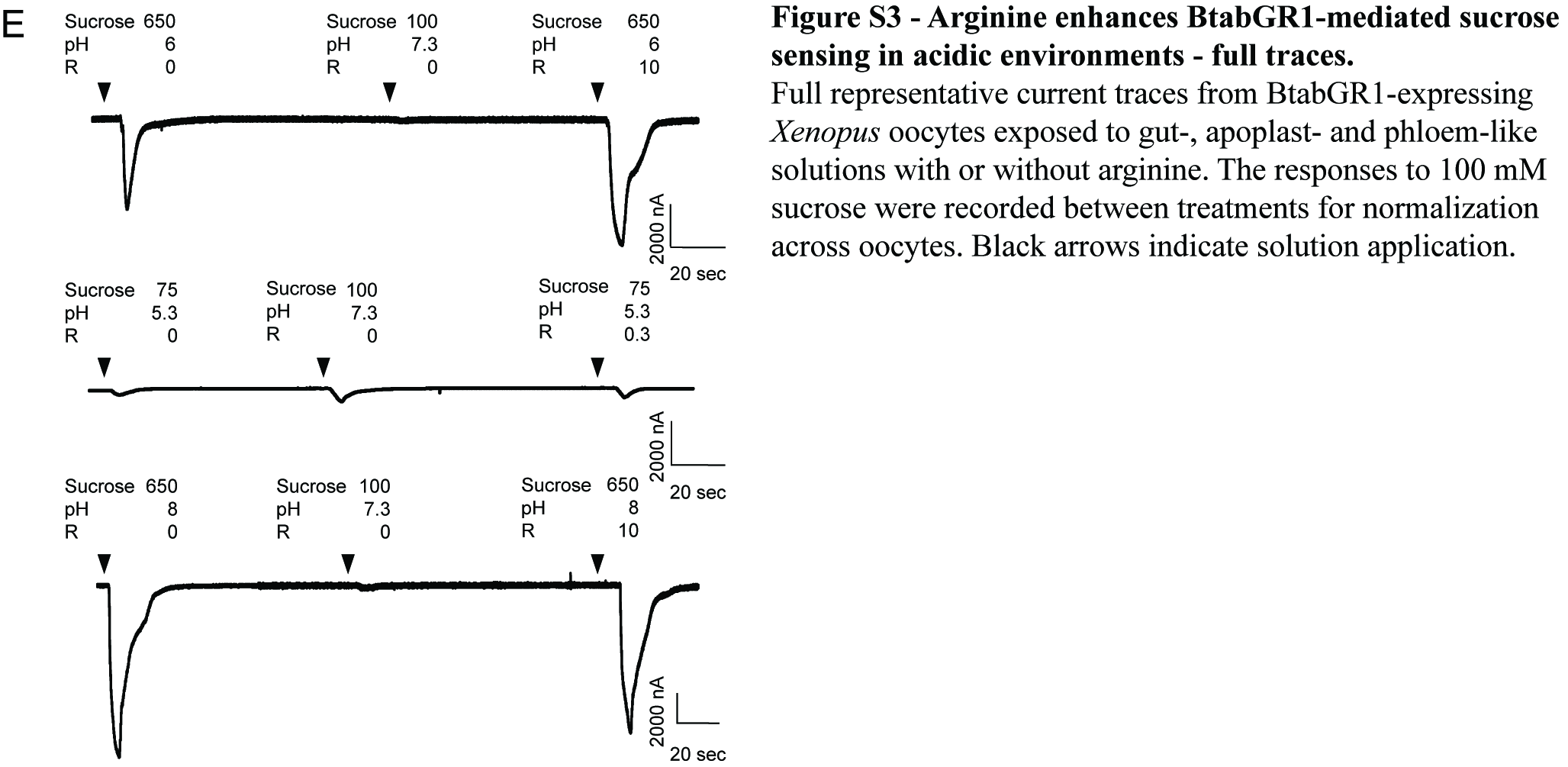

### Supplemental Figure 4

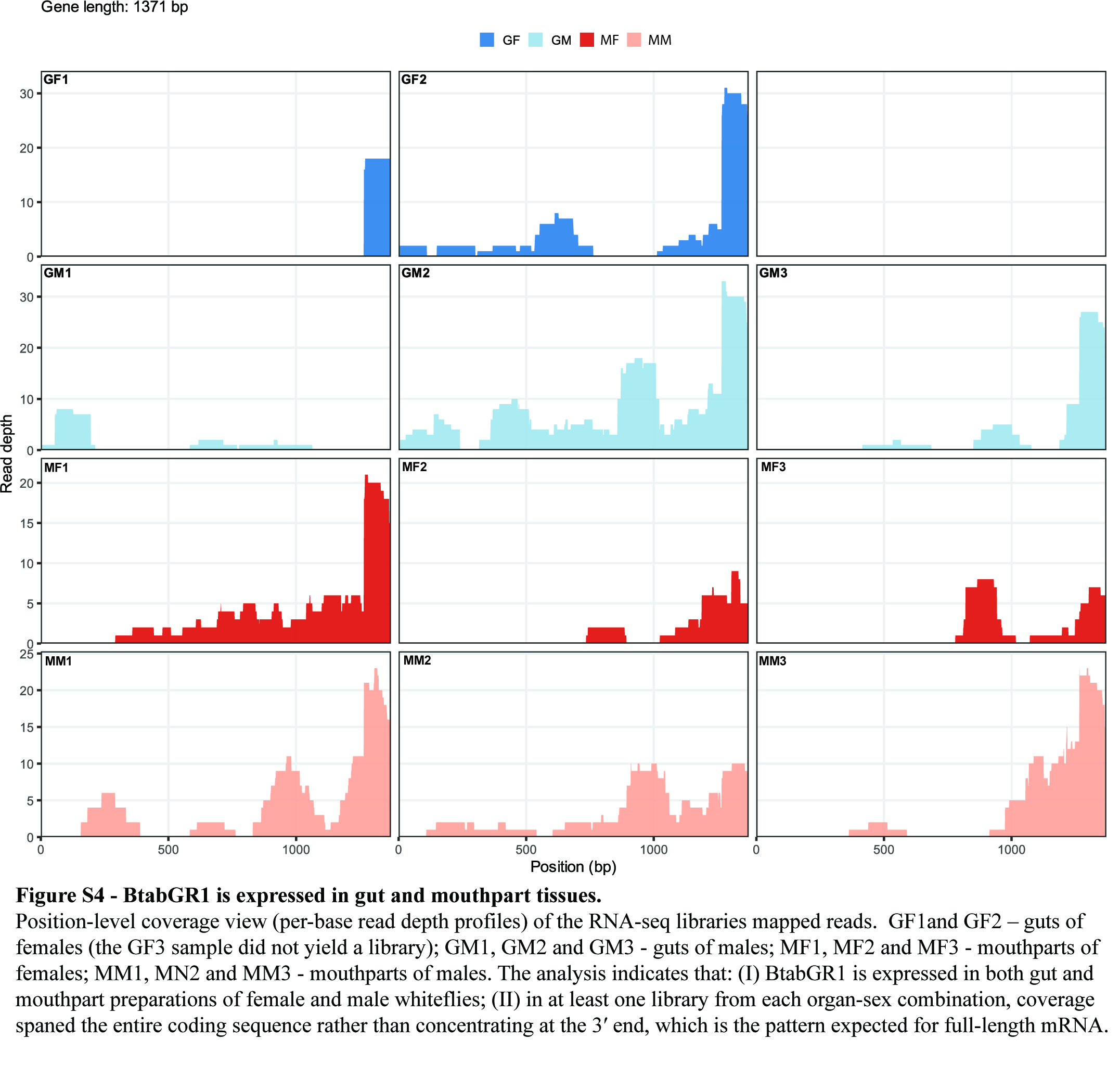
