## Supplemental Table 2 for "An Integrative Taste Receptor Links pH and Amino Acids to Sugar Sensing in *Bemisia tabaci*"

**Table S2. List of primers used in the qRT-PCR analysis presented in Figure 5B.**

| **#** | **Name** | **Sequence (5' -> 3')** |
| --- | --- | --- |
| 1 | qRT-PCR:Bta04282 (RPL13A) Forward | CATTCCACTACAGAGCTCCA |
| 2 | qRT-PCR:Bta04282 (RPL13A) Reverse | TTTCAGGTTTCGGATGGCTT |
| 3 | qRT-PCR:Bta10134 (sweet GR) Forward | GTGGACTTCGTGTTTGTGAC |
| 4 | qRT-PCR:Bta10134 (sweet GR) Reverse | CTCCGCCTTCTCCCATTG |
